# The cell and stress-specific canonical and non-canonical tRNA cleavage

**DOI:** 10.1101/2020.02.04.934695

**Authors:** Sherif Rashad, Teiji Tominaga, Kuniyasu Niizuma

## Abstract

Following stress, tRNA is cleaved to generate tRNA halves (tiRNAs). These stress-induced small RNAs have been shown to regulate translation during stress. To date, angiogenin is considered the main enzyme that cleaves tRNA at its anti-codon site to generate 35 ~ 45 nucleotide long 5′ and 3′ tiRNA halves, however recent reports indicate the presence of angiogenin-independent cleavage. We previously observed tRNA cleavage pattern occurring away from the anti-codon site. To explore this non-canonical cleavage, we analyze tRNA phenotypical cleavage patterns in rat model of ischemia reperfusion and in two rat cell lines. *In vivo* mitochondrial tRNAs were prone to this non-canonical cleavage pattern. *In vitro*, however, both cytosolic and mitochondrial tRNAs could be cleaved non-canonically. We also evaluated the roles of angiogenin and its inhibitor, RNH1, in regulating tRNA cleavage during stress. Our results suggest that mitochondrial stress has an important regulatory role in angiogenin-mediated tRNA cleavage. Angiogenin does not appear to regulate the non-canonical cleavage pattern of tRNA, and RNH1 does not affect it as well. Finally, we verified our previous findings of the stress-specific role of Alkbh1 in regulating tRNA cleavage and showed a strong influence of stress type on Alkbh1-mediated tRNA cleavage and that Alkbh1 impacts non-canonical tRNA cleavage.

## Introduction

Understanding the process of protein translation is indispensable for our understanding of cellular physiology as well as pathology. How cells regulate translation during adverse stress is key in our understanding of cellular stress pathways and death mechanisms. Over the past decades, our view of tRNA has changed from a static element in the translation complex into a dynamic player in the process of translation regulation^1^. During stress, cells can change tRNA transcripts abundance to drive the translation towards the generation of antioxidant proteins^2^. The same process was shown to drive metastasis in breast cancer^3^. tRNA modifications play an important role in regulating transcript availability and efficient protein translation^4,5^. tRNA is also a source of small non-coding RNAs that have an important regulatory role in translation, namely: tiRNAs and tRFs (tRNA derived fragments produced via DICER)^6,7^.

tiRNAs are small non-coding tRNA derived fragments that are produced via tRNA cleavage during stress and under physiologic conditions, although their production exponentially increases during stress^6,8^. tiRNAs are currently under the scope to be used as biomarkers for diseases such as stroke and malignancies^9–11^, nonetheless, the process of tRNA cleavage is not well understood in terms of regulation and exact function^6^. Canonically, tRNAs are cleaved by Angiogenin (Ang) at the anti-codon site to generate tRNA halves (tiRNAs)^12,13^, however, recent studies have identified Ang-independent cleavage^14–16^. In addition, we observed in our previous report the existence of larger 5’tiRNA fragments implying cleavage away from the anti-codon site (i.e. non-canonical cleavage)^17^. Moreover, does these small RNAs play a role beyond the reported protein translation repression or not is still not known^6,7,18,19^. The cleavage process of tRNA was shown to be regulated, at least at some level, via tRNA methylation modifications such as 1-methyadenosine (m^1^A) modifications and 5-methylcytosine (m^5^C) modifications via a variety of tRNA methyltransferases or demethylases^4,20–23^. We recently reported that Alkbh1, a cytosolic and mitochondrial tRNA m^1^A demethylase, regulates cellular stress responses and tRNA cleavage in a stress specific manner^23^. This indicates a higher-level regulation rather than a global cleavage pattern, since not all tRNA species are equally modified and the affinity of certain tRNA methylation modifying enzymes is selective to particular tRNAs^4,20,21^.

While Ang-independent cleavage was reported *in vitro* and *ex vivo*^15,16^, it is unclear whether this process occurs *in vivo*. It is also unclear what enzymes or factors impact Ang-independent tRNA cleavage, what are the sites of this Ang-independent cleavage and how this process is related to cellular stress response. To address these questions, in particular the first one, we employed an animal model of cerebral ischemia; transient middle cerebral artery (tMCAO) ischemia-reperfusion model (I/R)^24^. We utilized Northern blotting to evaluate cleavage site of various cytosolic and mitochondrial tRNAs after cerebral ischemia. We also evaluated the same process phenotypically in 2 rodent neuronal cell lines, B35 neuroblastoma and PC12 pheochromocytoma. Using these combined approaches, we were able to observe non-canonical tRNA cleavage (away from the anti-codon) *in vivo* and *in vitro*. We also show that Ang does not impact this non-canonical cleavage especially in mitochondrial tRNAs, indicating the potential role of other unidentified tRNA RNAse(s), while show that m^1^A demethylation by Alkbh1 does enhance the mitochondrial tRNA non-canonical cleavage.

## Results

### Mitochondrial tRNAs are cleaved non-canonically *in vivo*

Previously, we detected that several tRNA species were cleaved into multiple fragments, with some of these fragments having sizes essentially indicating that their cleavage was not at the canonical anti-codon site^23^. Moreover, these non-canonical fragments were regulated, at least in part, by the m^1^A demethylating action of Alkbh1^23^. Based on these results, we were interested in evaluating the nature of these fragments, how they correlate to stress and whether they occur also *in vivo* or only *in vitro*. Since these fragments were larger 5’tiRNAs, indicating cleavage towards the heavily modified 3’ side of the tRNA, we expected heavy biases from available sequencing technologies that would hinder the elucidation of these fragments. To that end, we utilized a rat model of cerebral ischemia, the tMCAO model ^9^, which yielded enough RNA allowing us to examine cytosolic and mitochondrial tRNA cleavage using Northern blotting to test our hypothesis. We, and others, have shown that tMCAO model induces tRNA cleavage^9,25^. We have reported previously in the same model that the maximum tRNA cleavage is observable 24 hours after reperfusion^9^. Thus, we selected this time point for our evaluation using DIG-labeled DNA probes based Northern blotting. Here, we wanted to evaluate whether tRNA cleavage after I/R injury is a global phenomenon or specific to particular tRNA-species as in tGCI and whether we will observe the non-canonical cleavage in mt-tRNAs as well.

Following I/R, the cleavage pattern of cytosolic tRNAs was selective to particular species (Figure 1A). tRNA-Arg^CCT^ was not cleaved *in vivo*. Also, tRNA-Met^CAT^ (also known as initiator tRNA-Met or tRNA^iMET^) was not cleaved after I/R. While tRNA-Gly^GCC^, tRNA-Leu^AAG^ and tRNA-Ala^AGC^ were robustly cleaved. tRNA-Gln^CTG^ was only minimally cleaved, and the membrane needed extended exposure periods in order to reveal the 5’tiRNA fragments resulting from this cleavage. The size of the cleaved cytosolic 5’tiRNAs indicated a canonical codon cleavage, given that cytosolic cleavage. However, Leu^AAG^ had a larger non-canonical 5’tiRNA fragment that was present in Sham as well as I/R samples.

**Figure 1:**
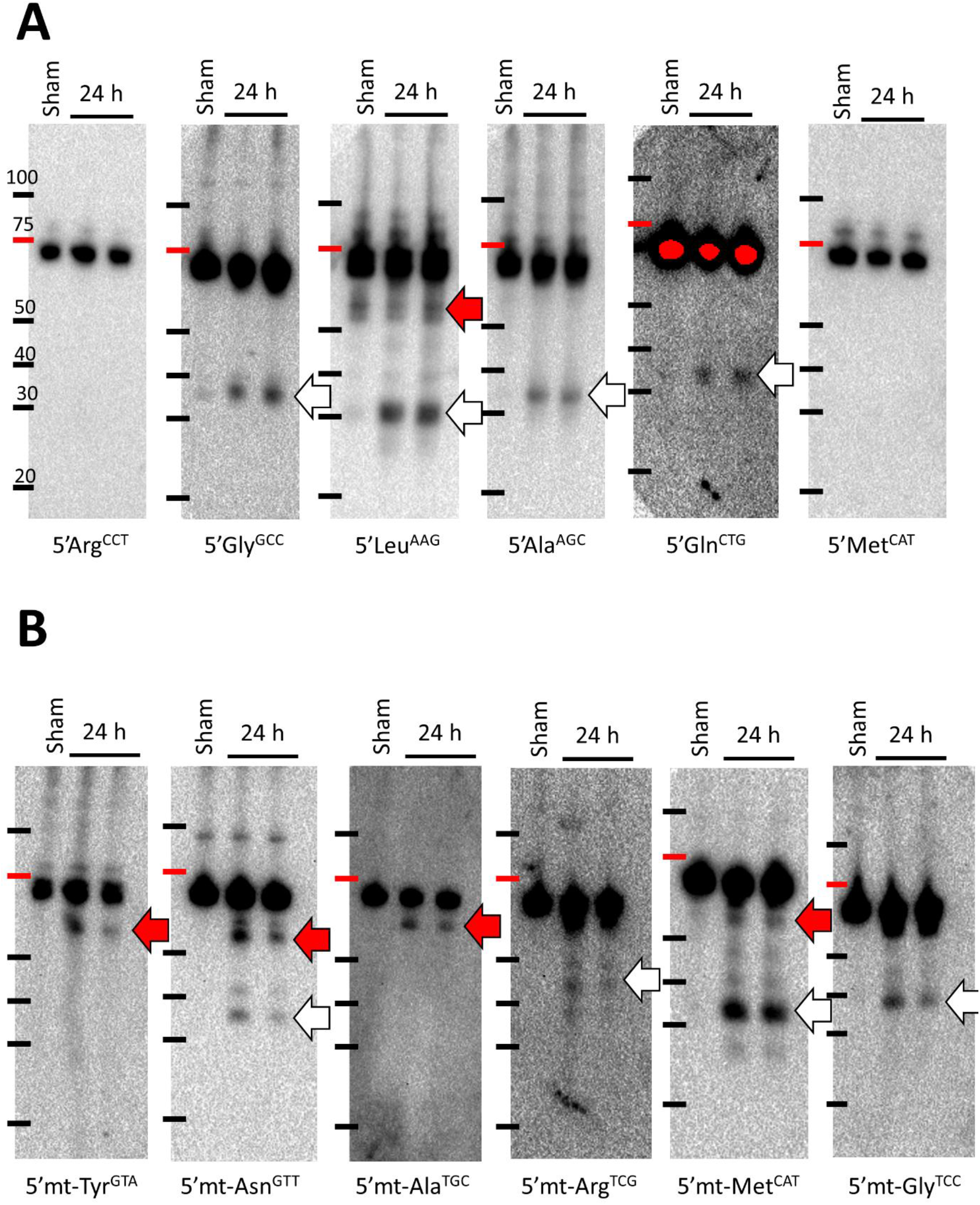
Analysis of cytosolic (A) and mitochondrial (B) tRNA cleavage after ischemia-reperfusion injury in rats after 24 hours of reperfusion. 4μg total RNA were used per lane (sample). RNA concentration was confirmed immediately prior to electrophoresis. **A:** Cytosolic tRNAs and **B:** Mitochondrial tRNAs

Mitochondrial tRNAs (Figure 1B) were also cleaved, as evident from the analysis of several mt-tRNAs. Interestingly, the phenotypical appearance of the cleavage pattern appears to deviate from the canonical angiogenin-mediated tRNA cleavage visible with cytosolic tRNA species in some of the mitochondrial tRNA species, as observed in the tGCI model. While the 5’tiRNA fragments resulting from cytosolic tRNA cleavage were typically between the 35 to 45 nucleotide mark, indicating canonical cleavage at the anti-codon, the 5’tiRNA fragment from mt-Tyr^GTA^, mt-Ala^TGC^ and some mt-Asn^GTT^ fragments were much larger, being detected between 50 and 60 nucleotides mark. mt-Asn^GTT^ GTT cleavage was bimodal, with fragments at the canonical size of 35 to 45 nucleotides and other non-canonical larger fragments similar to what was observed with mt-Tyr^GTA^. mt-Met^CAT^ and mt-Gly^TCC^ were largely canonically cleaved, although mt-Met^CAT^ exhibited several larger non-canonical 5’ fragments as well indicating multiple points of cleavage, similar to what we observed in a previous report^23^. Moreover, the near canonical size of mt-tRNAs does not rule out off anti-codon cleavage (i.e. non-canonical cleavage), which cannot be resolved using Northern blotting.

Collectively, these findings indicate that the process of *in vivo* cleavage of tRNA is vastly different between cytosolic and mitochondrial tRNAs, at least for the tested tRNAs. It also clearly indicates the presence of other cleavage sites (other than the anti-codon) for the generation of stress induced tiRNAs.

### *In vitro* tRNA cleavage is cell line and stress specific and shows non-canonical fragments

Next, we wanted to qualitatively evaluate the cleavage pattern of tRNA *in vitro*. In particular, we wanted to evaluate whether tRNA can be cleaved non-canonically *in vitro* and whether the cleavage patterns are the same under different stress conditions and in different cell lines. We exposed PC12 and B35 cells (both rat neuronal cell lines) to oxidative stressors using toxic doses of sodium meta-Arsenite (As)^12^ and Antimycin A (AM), which targets mitochondrial respiratory complex III^26^. Exposure of PC12 cells to As led to induction of apoptosis and robust tRNA cleavage (Supplementary figure 1A), while exposure to 200μg/ml of antimycin A, a dose that is 20 times the reported to induce pyroptosis and inflammasome activation in other cell lines ^26^, did not have significant effect on cell viability or gross tRNA cleavage. On the other hand, both As and antimycin A induced cell death and tRNA cleavage in B35 cells as evident by Annexin V assay and SYBR gold staining (Supplementary figure 1B). Interestingly, despite having similar levels of apoptosis with As and AM, tRNA cleavage was significantly more robust after AM stress in B35 cells. This might reflect a strong influence of mitochondrial OXPHOS inhibition induces stress on tRNA cleavage that is not yet explored.

To examine this notion, we used DIG-labeled DNA probes to evaluate individual tRNAs’ cleavage in both cell lines following As and antimycin stresses. First, we examined cytosolic tRNA cleavage in PC12 and B35 cells after As or antimycin stresses (Figure 2). We observed several differences in the cleavage patterns of tRNA between PC12 and B35 cells.

**Figure 2:**
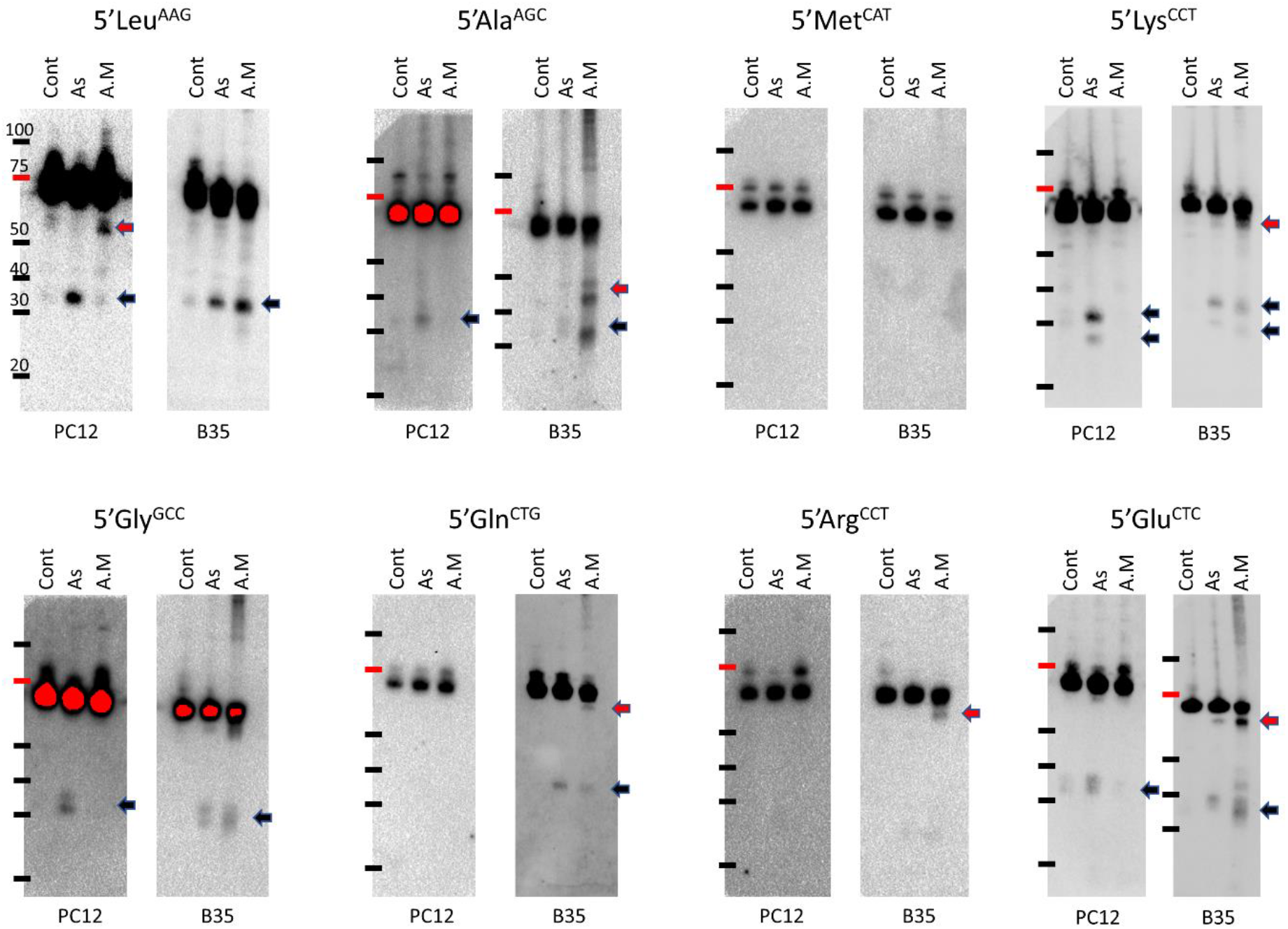
Analysis of cytosolic tRNA cleavage in B35 and PC12 cell lines after arsenite (As) or antimycin (A.M) stresses. Cells were exposed to As (1mM) or A.M (200μg/ml) for 4 hours (B35) or 6 hours (PC12). 3μg total RNA were loaded per lane (sample). Black arrow: Canonical fragments, red arrow: non-canonical fragments. Shorter exposure images are provided in supplementary figure 2 for Gly^GCC^ in B35 and PC12, Ala^AGC^ in PC12 and for Lys^CTT^ in B35.

Leu^AAG^ showed quite an interesting pattern of cleavage under different stresses and between the 2 cell lines. While As stress caused canonical cleavage of Leu^AAG^ in both cell lines, antimycin stress caused an enrichment of a non-canonical 5’half in PC12 cells with minimal enrichment of the canonical 5’halves. On the contrary, antimycin stress led to canonical cleavage of Leu^AAG^ in B35. Considering that PC12 cells were resistant to antimycin stress, there is an apparent correlation between the non-canonical cleavage of Leu^AAG^ and cellular responses to ischemia. Such a correlation needs further analysis, indeed, to evaluate if there is a causative link between this process and cellular fate.

Ala^AGC^ was only cleaved after As stress in PC12 canonically, but not after AM stress. It was cleaved both canonically and non-canonically after AM stress in B35 with minimal cleavage after As stress in B35 cells.

Gly^GCC^ was cleaved canonically in both cell lines, although antimycin stress in PC12 cells did not lead to robust cleavage. Gln^CTG^ was only cleaved in B35 cells into a canonical 5’tiRNA fragment after both stresses and an extra non-canonical fragment only after antimycin stress. Met^CAT^ was not cleaved in both cell lines. Arg^CCT^ was cleaved non-canonically after antimycin stress in B35 cells only resulting in a large 5’tiRNA fragment. Lys^CTT^ was cleaved into 2 5’tiRNA fragments after As stress in PC12 (and minimally after AM stress) and after As and antimycin stresses in B35. Both fragments were within the canonical range of tiRNAs (i.e 35 ~ 45 nt) suggesting that the cleavage occurred at the anticodon canonically and at a nearby site non-canonically. Moreover, Lys^CTT^ showed a large non-canonical tiRNA fragment after antimycin stress in B35 cells. Glu^CTC^ was also cleaved canonically after As stress in PC12 and after As and antimycin stresses in B35 with a non-canonical tiRNA fragment observed after both stresses in B35, albeit more evident after antimycin stress.

Mitochondrial tRNA cleavage also showed observable differences between PC12 and B35 cell lines (Figure 3). mt-Tyr^GTA^ was cleaved after As stress in PC12 (in a more or less degradation-like manner with many fragments observable) while it was cleaved canonically and non-canonically after antimycin stress in B35 but not As stress. mt-Met^CAT^ showed non-canonical cleavage after As and antimycin stresses in PC12 cells, although to a higher degree with As stress, while no observable cleavage was noted in B35 cells. mt-Asn^GTT^, mt-Gly^TCC^, mt-Ala^TGC^ and mt-Arg^TCG^ were not cleaved in both cell lines.

**Figure 3:**
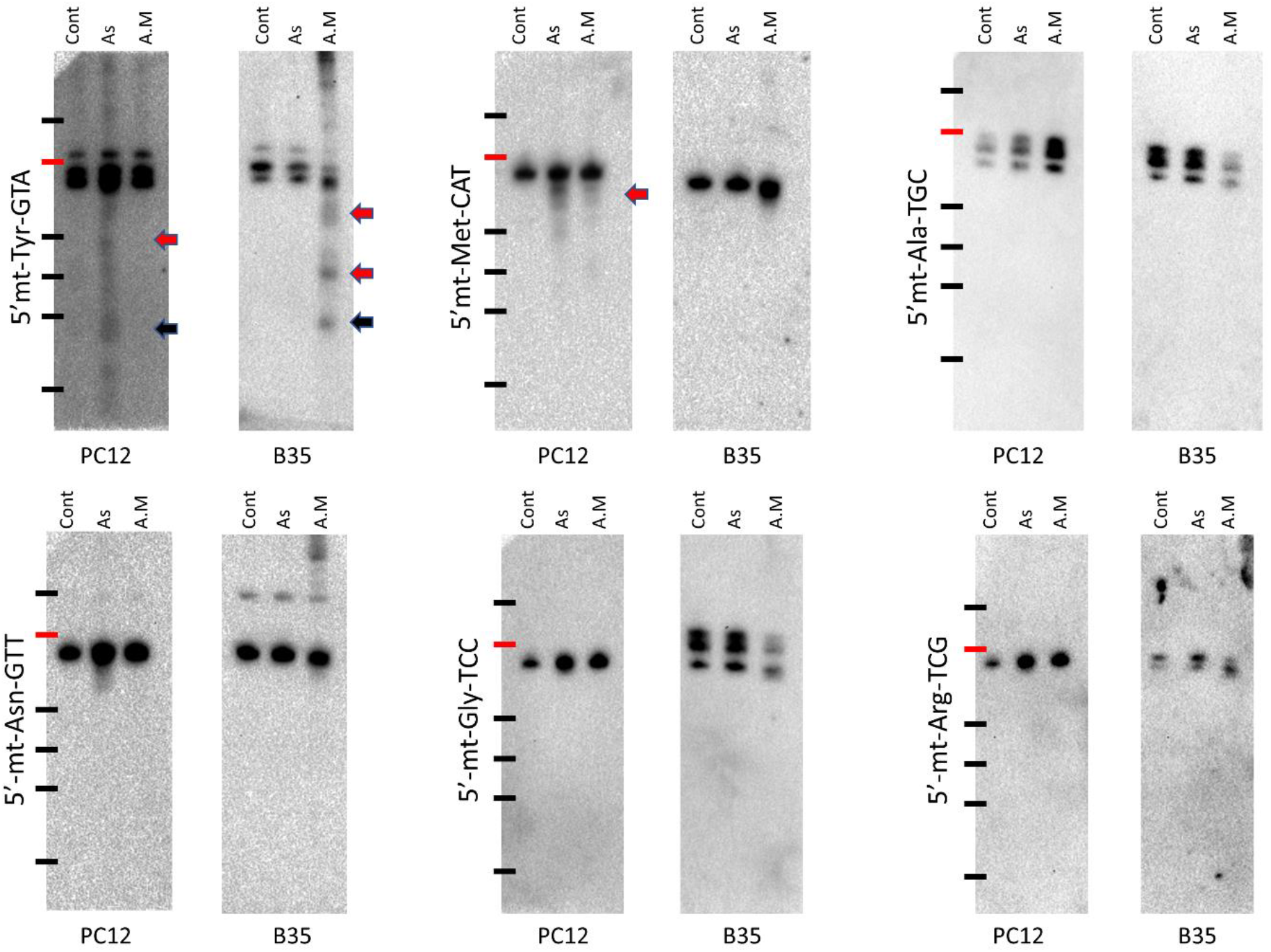
Analysis of mitochondrial tRNA cleavage in B35 and PC12 cell lines after arsenite (As) or antimycin (A.M) stresses. Cells were exposed to As (1mM) or A.M (200μg/ml) for 4 hours (B35) or 6 hours (PC12). 3μg total RNA were loaded per lane (sample). Black arrow: Canonical fragments, red arrow: non-canonical fragments.

It is notable to observe the differences exist not only between these cell lines under the same stress, but also between *in vivo* and *in vitro* patterns of tRNA cleavage and the tRNA species cleaved. Collectively, these results indicate a stress, cell type and tRNA specific pattern of cleavage that is drastically different between *in vivo* and *in vitro* systems and largely dependent on cellular stress mechanism. These results also show that non-canonical cleavage is specific and not a result of haphazard degradation of tRNA, since the same tRNA species were non-canonically cleaved in certain conditions and not the others.

Overall, we observed non-canonical cleavage *in vivo* to occur in mitochondrial tRNAs and *in vitro* to be more evident with the mitochondrial targeting stressor antimycin. This suggests a big role played in mitochondrial stress and dynamics in regulating canonical versus non-canonical tRNA cleavage.

### Ang and RNH1 overexpression; impact on tRNA cleavage after stress

Angiogenin is currently regarded as the main enzyme involved in tRNA cleavage^7,13,18,19,27^. RNH1 is the angiogenin inhibitor^13,28–31^, and its knockout was reported to increase cytotoxicity via what believed to be unchecked Angiogenin induced tRNA cleavage^28^. Recently, several reports have demonstrated Angiogenin-independent tRNA cleavage^15,16^ Here we sought to answer two questions. First, is Angiogenin responsible for the non-canonical tRNA cleavage we observed? And if not, what is the enzyme or possible enzymes responsible for this process? The second question was how specific is the Angiogenin mediated tRNA cleavage, and does Angiogenin cleave all tRNAs?

To answer these questions, we introduced DNA plasmids to overexpress human Angiogenin (hAng), as we reported previously^12^, or RNH1 into PC12 cells (Supplementary figure 3A and B). Then we exposed the cells to As (400μM) or Antimycin (200μg/ml) stresses for 4 hours and observed the impact of hAng or RNH1 overexpression on gross tRNA cleavage as well as species-specific tRNA cleavage.

First, hAng overexpression in PC12 cells increased gross tRNA cleavage following As stress but not antimycin stress as observed by SYBR gold staining (Supplementary figure 3C). RNH1 overexpression, however, did not lead to the expected effect of suppressing gross tRNA cleavage (Supplementary figure 3D).

To further characterize these findings, we looked into several tRNA species’ specific cleavage patterns following gene overexpression. Leu^AAG^, which showed canonical and non-canonical 5’tiRNA fragments especially after antimycin stress, showed an interesting pattern. While hAng enhanced the canonical Leu^AAG^ 5’tiRNA fragments especially after antimycin stress, it had no effect on the larger non-canonical fragments. Moreover, RNH1 overexpression reduced the canonical 5’tiRNA fragments following antimycin stress and not after As stress, and again had no impact on the non-canonical fragments (Figure 4).

**Figure 4:**
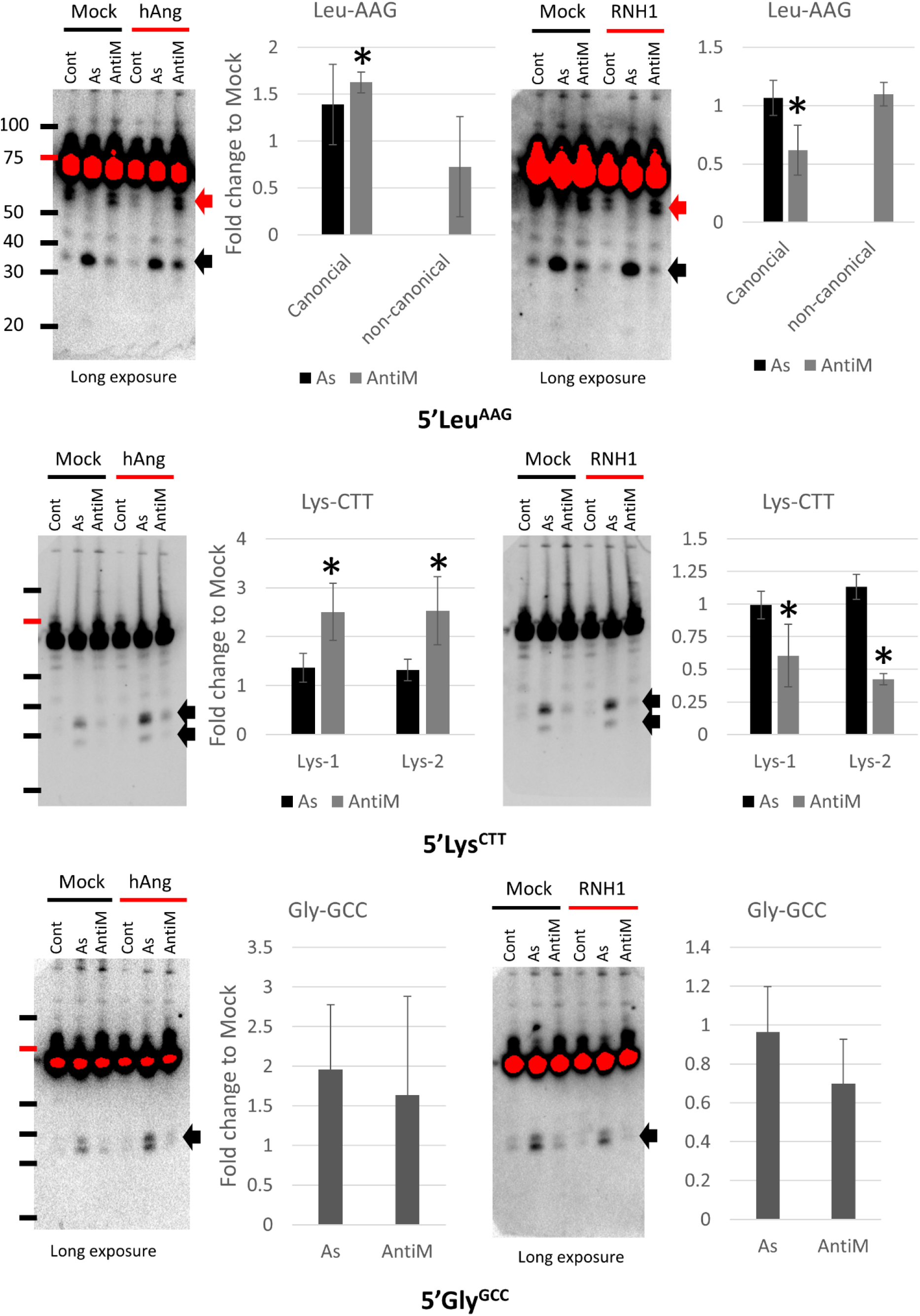
Impact of hANg or RNH1 overexpression on stress induced tRNA cleavage. Specific tRNAs were probed using DIG-labeled probes to evaluate the impact of hANg or its inhibition by RNH1 on their cleavage in PC12 cells following arsenite (As) or antimycin A (AntiM) stresses. Black arrow: canonical fragments, red arrow: non-canonical fragments. Shorter exposure images for the overexposed blots are presented in supplementary figure 4. 5s rRNA was used as a loading control and presented in supplementary figure 4. Graphs represent quantification from 3 independent experiments. Asterisk: statistically significant (*p* < 0.05) and fold change > 1.5 folds.

hAng enhanced the cleavage of Lys^CCT^, which was cleaved in 2 fragments with sizes suggesting cleavage at or near the anti-codon, after AM stress but not after As stress, and RNH1 reversed this pattern (Figure 4). hAng enhanced Ala^AGC^ cleavage after both As and AM stresses, RNH1, on the other hand, only reversed this pattern with AM stress and not with As stress (Figure 5).

**Figure 5:**
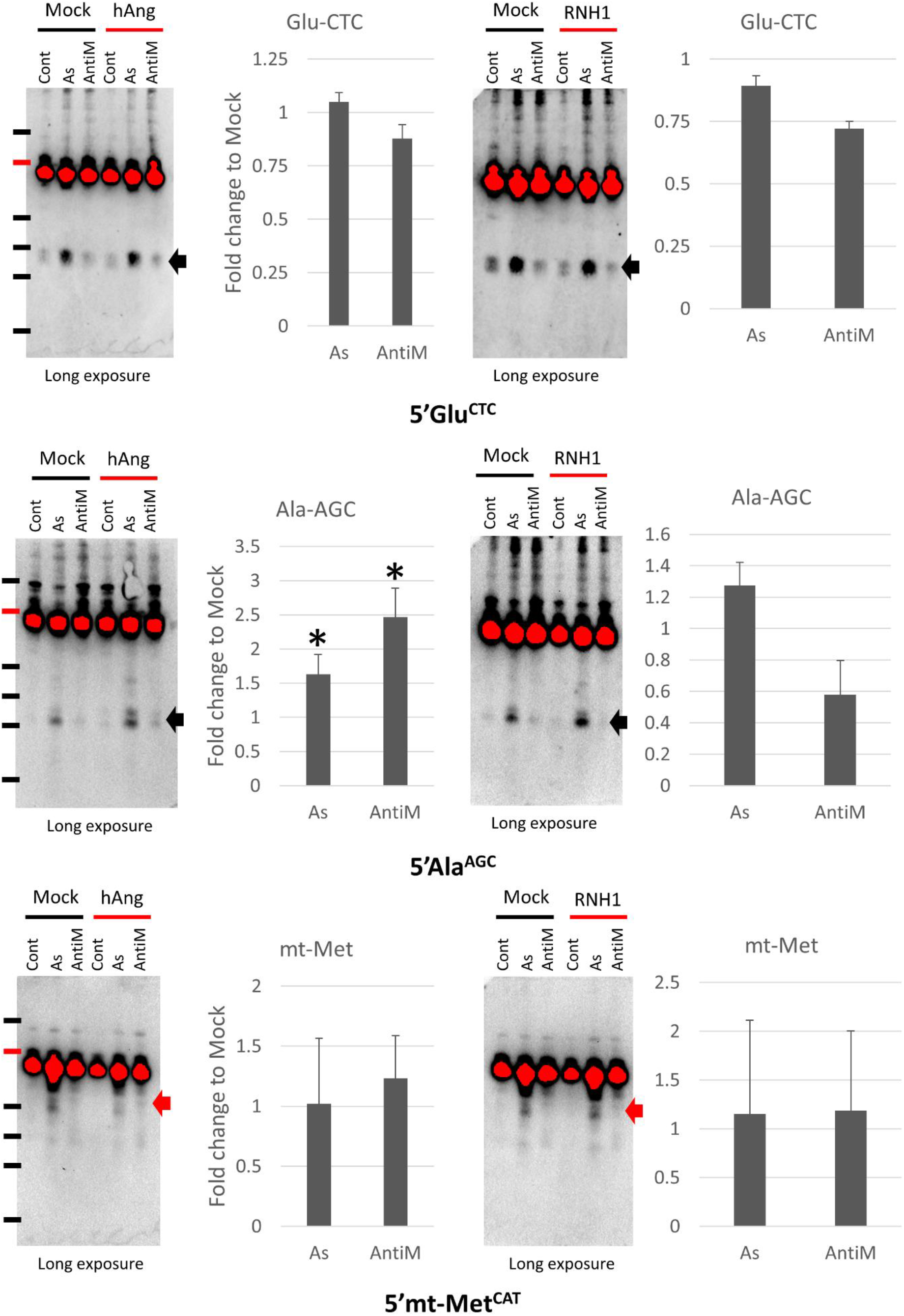
Impact of hANg or RNH1 overexpression on stress induced tRNA cleavage (continued). Black arrow: canonical fragments, red arrow: non-canonical fragments. Shorter exposure images for the overexposed blots are presented in supplementary figure 4. 5s rRNA was used as a loading control and presented in supplementary figure 4. Graphs represent quantification from 3 independent experiments. Asterisk: statistically significant (*p* < 0.05) and fold change > 1.5 folds.

Glu^CTC^ (Figure 4) and Gly^GCC^ (Figure 5) did not exhibit significant changes in the levels of their 5’tiRNA fragments after hAng or RNH1. mt-Met^CAT^, which is cleaved mainly non-canonically, was not affected also by hAng or RNH1, indicating a possible independence of mitochondrial tRNA cleavage from Ang.

These results indicate that the impact of angiogenin on tRNA cleavage is not only stress specific, but it is also limited to specific tRNA species^15^. Moreover, it indicates that angiogenin has no impact on the non-canonical 5’tiRNA fragments generation, whether cytosolic or mitochondrial, at least in the studied transcripts.

### Impact of Alkbh1-mediated m^1^A demethylation on tRNA cleavage

Previously, we examined the role of Alkbh1 on tRNA^23^. Alkbh1 is a 1-methyladenosine (m^1^A) demethylase and a histone dioxygenase^4,32^. We showed that Alkbh1 regulates tRNA cleavage and cellular fate under stress in a specific pattern in B35 cells^23^. We also observed that Alkbh1 impact on tRNA cleavage can be variable between canonical and non-canonical fragments^23^. Therefore, we examined the impact of Alkbh1 on the non-canonical cleavage pattern we observed herein in PC12 cells using Alkbh1 overexpression plasmid [Supplementary figure 5]. Alkbh1 overexpression had a stress specific impact on tRNA cleavage similar to what we observed previously^23^, with As stress causing more robust upregulation of tiRNAs after Alkbh1 overexpression as compared to AM stress [Figure 6]. Paradoxically, As stress did not affect the non-canonical mt-Met^CAT^ fragments after Alkbh1 overexpression, while AM stress greatly enhanced the cleavage of this mitochondrial tRNA, further signifying the role of mitochondrial stress in regulating tRNA cleavage. On the other hand, the non-canonical Leu^AAG^ fragment was not impacted by Alkbh1. This signifies a complex regulatory relation between specific tRNA m^1^A methylation and cleavage patterns.

**Figure 6:**
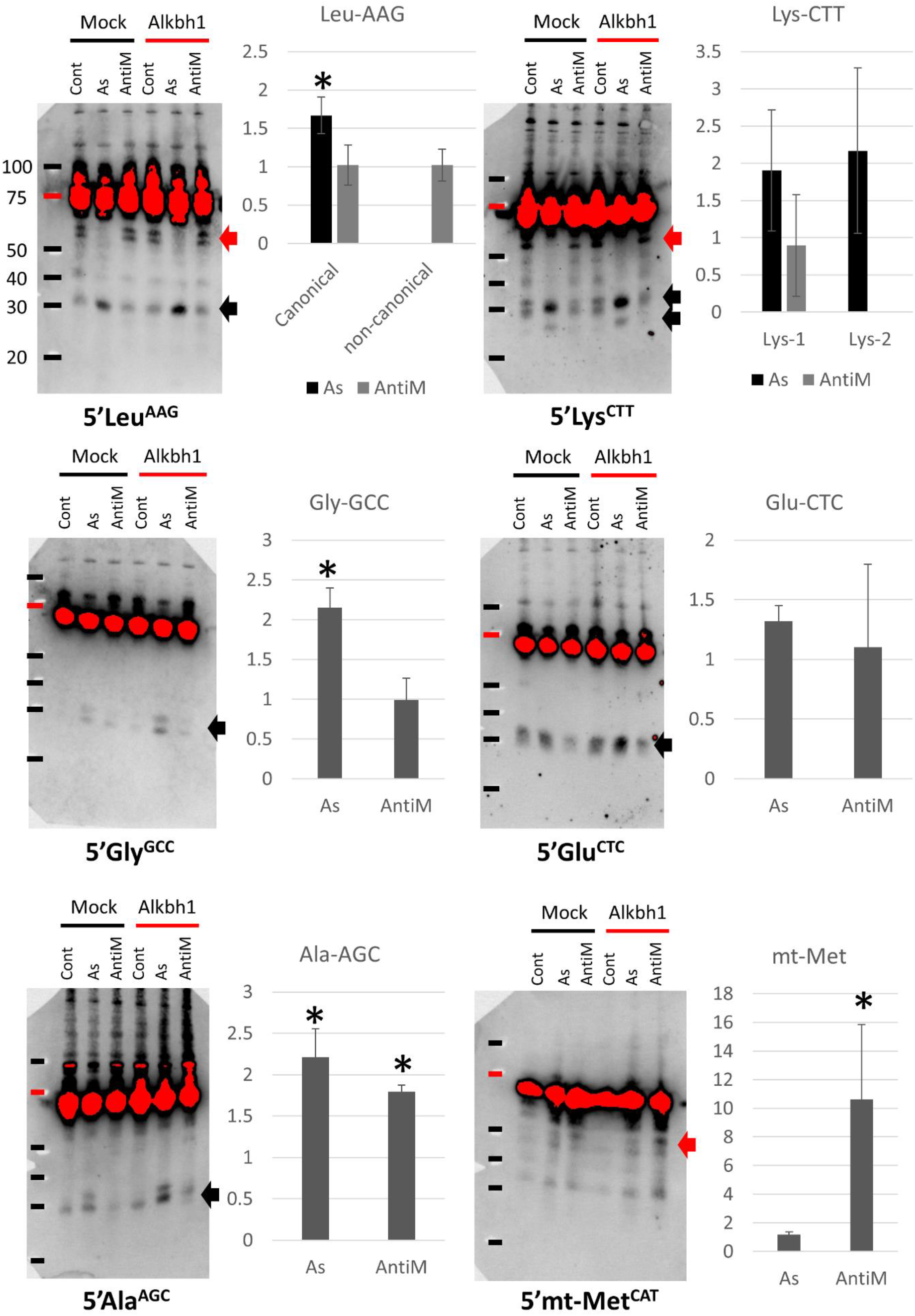
Impact of Alkbh1 overexpression on stress induced tRNA cleavage. Black arrow: canonical fragments, red arrow: non-canonical fragments. Shorter exposure images for the overexposed blots are presented in supplementary figure 6. 5s rRNA was used as a loading control and presented in supplementary figure 6. Graphs represent quantification from 3 independent experiments. Asterisk: statistically significant (*p* < 0.05) and fold change > 1.5 folds.

## Discussion

In this report, we show for the first time, to the best of our knowledge, that certain tRNA transcripts can be cleaved at multiple cleavage points, and that the cleavage is not restricted to anti-codon site, to generate varying lengths of tiRNAs. This non-canonical cleavage occurs *in vivo* mainly in mitochondrial tRNAs, while *in vitro* it can affect both cytosolic and mitochondrial transcripts leading to the generation of multiple large tiRNAs with sizes depending on the tRNA transcript cleaved. Angiogenin independent cleavage was reported previously by Su et al^15^, however, in their report they did not reveal the non-canonical fragments revealed herein. This might be the fact that when the cleavage site shifts towards 3’ side of tRNAs, the abundant modifications in that part of tRNA introduce biases in the sequencing hindering the resolution of the cleavage sites and the proper sequencing of the larger fragments^33^. Notwithstanding, we became aware of new sequencing methodologies that can solve these issues^33^. However, these methods were not available during the preparation of this work.

The major limitation of not sequencing the larger non-canonical fragments is the inability to resolve their functional roles using bioinformatics or biological approaches. Moreover, since their sequences are similar in a large part to their canonical counterparts, it will be difficult and laborious to isolate them from biological samples and perform functional studies^34^. Therefore, it is imperative that future work incorporates advanced sequencing technologies to resolve the nature of these non-canonical fragments and the role they play in regulating stress response.

It seems, however, that mitochondrial stress plays an important role in tRNA cleavage that was not explored in the literature. Previously, we reported that antimycin A, the complex III inhibitor, promoted mt-Met^CAT^ cleavage in Alkbh1 overexpressing cells without an apparent impact on cell fate^23^. The same dissociative relationship between tRNA cleavage and cell fate was observed in this work, where antimycin A caused a more robust gross tRNA cleavage compared to arsenite stress, despite having equal levels of apoptotic cells in B35 cells. Both arsenite and antimycin stresses promoted the increased certain tRNA transcripts cleavage after hAng overexpression in PC12 cells, on the contrary, overexpressing RNH1 seemed to reduce tRNA cleavage with antimycin stress only. RNH1 can be inactivated with general oxidants or with thiol-specific oxidants, rendering it inactive^35^. Indeed, arsenite is a general oxidant, which can explain the absence of response in RNH1 overexpressing cells exposed to arsenite in terms of tRNA cleavage. However, antimycin A, as a mitochondrial specific stress, may leave RNH1 unscathed, which can be considered as an important example of oxidative stress compartmentalization in regulating tRNA cleavage and the resulting translational effects. This theory, however, remains to be confirmed experimentally.

It is important to note that these non-canonical fragments are not products of non-specific cleavage or degradation of tRNAs. This is evident from the impact of various enzymes studied here on cytosolic versus mitochondrial non-canonical fragments. If these fragments were the result of non-specific cleavage, we would not have observed the stress specific/transcript specific and enzyme regulated changes in their level. How these fragments are generated, what’s their biological function and what are the regulatory mechanisms behind this process are still open questions that need to be addressed.

In conclusion, our presented work, in tandem with our previous report^23^ show that tRNA can be cleaved at multiple sites to generate non-canonical tiRNAs. These non-canonical tiRNAs are still not well understood, and future work should aim to identify their functional role, if any, using new sequencing methods. Our work also shows the impact of stress type on tRNA cleavage (for example, general oxidative stress versus mitochondrial specific oxidative stress). This is not yet explored as most reports on tRNA cleavage has used general oxidative stressors (such as H2O2 and arsenite). In addition, our work shows, for the first time, large differences between *in vivo* and *in vitro* tRNA cleavage patterns as well as differences between cell lines. These differences, in essence, will have an impact on our understanding of various disease models. This indicates the need for full characterization of tRNA cleavage under various stress conditions and in different cell lines and animal models in order to understand the full scope of this process.

## Methods

### Animals

transient middle cerebral artery occlusion (tMCAO) was induced in 8 weeks old, 180-200gm male Wistar rats as previously described^24^. Animals were bought from a local vendor (Kumagai-Shoten Sendai) and housed in a controlled environment with 12 hours light/dark cycle and temperature regulated at 23°C. Animals were allowed to acclimatize for at least 24 hours before any experimental procedures. All experiments were conducted according to protocols approved by the animal care facility of Tohoku University (Ethical approval number 2017MdA-137-3) and according to the ARRIVE (Animal Research: Reporting In Vivo Experiments) guidelines. All animal surgeries were performed between 9am and 4pm. No blinding was performed for this work.

### Animal model for transient middle cerebral artery occlusion (tMCAO)

The ischemia-reperfusion model was performed as described previously^24^. In brief, after induction of anesthesia using inhalation anesthesia by 66% nitrous oxide and 33% oxygen plus 1.5% isoflurane gas mixture, a midline incision was performed, carotid bifurcation exposed, external carotid artery ligated and incised and a coated occlusion suture (4035PKRe5, Doccol, Corp., Sharon, MA, http://www.doccol.com/) was inserted to occlude the middle cerebral artery for 60 min. After the occlusion period the suture was removed to induce reperfusion injury, the wound sutured, and the animals allowed to rest before being sacrifice at the designated time points based on a simple randomization scheme designed in Excel (Microsoft corp.). after reperfusion. Animals from were sacrificed after 24 hours. Sham animals were prepared by performing all the surgical steps without suture occlusion of the MCA.

### Cell culture and *in vitro* stress

2 cell lines were used in this study. PC12 (rat pheochromocytoma cell line, ATCC, Cat# CRL-1721) and B35 (rat neuroblastoma cell line, ATCC, Cat# CRL-2754). PC12 was cultured in RPMI 1640 medium (Wako, Osaka, Japan. Cat# 189-02025) supplemented with 10% heat inactivated horse serum (Gibco, Japan. Cat# 16050-122) and 5% fetal bovine serum (FBS, Hyclone, GE-life sciences, USA. Cat# SH30910.03). B35 cells were cultured in high glucose DMEM medium (Nacalai Tesque, Cat# 08457-55) supplemented with 10% FBS. No antibiotics were added to the growth media. Cellular stress was performed by adding sodium meta-Arsenite (Sigma-Aldrich, Cat# S7400-100G) or streptococcal antimycin A (Sigma-Aldrich, Cat# A8674) at the designated concentrations and durations to the cells in full growth medium.

### Plasmid transfection

1 × 10^6^ cells (PC12 cell line) were cultured per well in a 6 well plate coated by Poly-L-Lysine. FLAG-tagged human angiogenin plasmid (Cat# RC208874; OriGene; Rockville, MD), GFP-tagged rat RNH1 plasmid (Cat# ORa42809C, Genescript; Tokyo, Japan), GFP-tagged rat Alkbh1 plasmid (Cat# ORa11821, Genescrip; Tokyo, Japan) or Mock plasmid were transfected to the cells using Lipofectamine 3000 (Thermo Fischer, Cat# L3000015) as per manufacturer’s protocol in full growth medium. Three days after transfection the cells were evaluated for transfection efficiency using western blotting or qPCR. Cell stress was performed 72 hours after transfection as per experimental protocol.

### RNA isolation and extraction

At the designated time-points animals were euthanized using overdose isoflurane anesthesia and then transcardiac perfusion using ice cold saline was performed. Brains were extracted and sliced on a matrix and the middle third of the left cerebral hemisphere was lysed in Qiazol (Qiagen, Hilden, Germany, Catalog# 79306). RNA was extracted using the miRNeasy micro kit (Qiagen, Hilden, Germany, Catalog# 217084) with a DNase digestion step as per manufacturer’s protocol. RNA concentration was analyzed using nanodrop one (Thermo Fisher Scientific, San Jose, CA, Catalog# ND-ONE-W) prior to each experiment.

### Northern blot and SYBR gold staining

tiRNA generation using SYBR gold staining was performed as we previously described^10,12^. In brief, 4μg of total RNA were loaded per well after denaturation by heating at 70°C for 3 minutes in sample buffer (Novex TBE-Urea sample buffer; Thermo fisher Scientific, Catalog# LC6876). Samples were electrophoresed using Novex 15% TBE-Urea Gels (Invitrogen, Carlsbad, CA, Catalog# EC68755;) in 1x TBE buffer (Bio-Rad, Catalog# 161-0733). Gels were then incubated with 10,000-fold diluted SYBR Gold nucleic acid gel stain (Life technologies, Eugene, OR, Catalog# S11494) in 0.5× TBE buffer for 30 min at room temperature. Stained gels were visualized using Chemi-Doc MP Imager (Bio-Rad, Hercules, CA) and analyzed by Image Lab Software (Bio-Rad, Hercules, CA. RRID# SCR_014210).

### DIG-labeled probes based tiRNA detection

Detection of individual tRNA and tRNA species using targeted DIG-labeled probes was performed as we reported previously^10,12,17^. Following membrane transfer and cross linking, membranes were pre-hybridized in DIG Easy Hyb solution (Roche, Catalog# 11603558001) for 30 minutes at 40°C followed by overnight incubation at 40°C with 1 μg/ml DIG-labeled DNA probes in DIG Easy Hyb solution with continuous agitation. After two low stringency washes for 5 minutes each with 2× saline sodium citrate/ 0.1% sodium dodecyl sulfate buffer (2x SSC buffer; Promega Co., Madison, WI, Catalog# V4261) and a high stringency wash for 2 min with 1x SSC buffer, the membrane was blocked in blocking reagent (Roche, Catalog# 11585762001) for 30 min. The membranes were then probed with alkaline phosphatase labeled anti-DIG antibody (Roche, Catalog# 11093274910) for 30 min, washed for 30 min with washing buffer (1 × maleic acid buffer + 0.3% Tween 20; Roche, Catalog# 11585762001), then equilibrated for 3 min in detection buffer (Roche, Catalog# 11585762001). Signals were visualized with CDP-Star Ready-to-Use (Roche, Catalog# 12041677001) and images taken with Chemi-Doc MP Imager.

Probe sequences used in this study were:

For cytosolic tRNA:

5'tRNA-Arg^CCT^: 5'-AGGCCAGTGCCTTATCCATTAGGCCACTGGGGC

5'tRNA-Leu^AAG^: 5'-AATCCAGCGCCTTAGACCGCTCGGCCATGCTACC

5'tRNA-Gln^CTG^: 5'-AGTCCAGAGTGCTAACCATTACACCATGGAACC

5'tRNA-Gly^GCC^: 5’-GCAGGCGAGAATTCTACCACTGAACCACCAATGC

5'tRNA-Ala^AGC^: 5’-CTAAGCACGCGCTCTACCACTGAGCTACACCCCC

5’tRNA-Met^CAT^: 5’GACTGACGCGCTACCTACTGCGCTAACGAGGC

5’tRNA-Lys^CTT^: 5’GTCTCATGCTCTACCGACTGAGCTAGCCGGGC

5’tRNA-Glu^CTC^: 5’GCGCCGAATCCTAACCACTAGACCACCAGGGA

For mitochondrial tRNA:

5'mit-tRNA-Asn^GTT^: 5'-AGCTAAATACCCTACTTACTGGCTTCAATCTA

5'mit-tRNA-Tyr^GTA^: 5'-AGTCTAATGCTTACTCAGCCACCCCACC

5’mt-tRNA-Met^CAT^: 5’CCCGATAGCTTAGTTAGCTGACCTTACT

5’mt-tRNA-Arg^TCG^: 5’TCATTAATTTTATTTAAACTAATTACCA

5’mt-tRNA-Gly^TCC^: 5’GTCAGTTGTATTGTTTATACTAAGGGAGT

5’mt-tRNA-Ala^TGC^: 5’ATCAACTGCTTTAATTAAGCTAAATCCTC

Loading control: 5s rRNA: 5’5'AAAGCCTACAGCACCCGGTATTCCC

Sequences for Cytosolic tRNAs were acquired from GtRNAdb (http://gtrnadb.ucsc.edu/) and for mitochondrial tRNAs from mitotRNAdb (http://mttrna.bioinf.uni-leipzig.de/mtDataOutput/).

Following acquisition of images using the DIG-labeled probes, membranes were stripped using 0.1x SSC – 0.5% sodium dodecylsulphate (SDS) solution at 80°C for 30 minutes then washed briefly in PBS-T and re-probed or stored in 5x SSC buffer for long term storage at 4°C.

Quantification was performed using ImageJ software^36^. Data were presented as the ratio (fold change) between target gene overexpressing cells and Mock transfected cells per given stress. Statistical analysis (Student’s t-test) was conducted on SPSS v.20 (IBM Corp.). Statistical significance was considered when *p* was less than 0.5 and fold change of > 1.5 folds were observed.

### Western blotting

Following plasmid transfection, cells were collected and lysed in NE-PER nuclear and cytoplasmic extraction reagent (Thermo Fischer, Cat# 78833). The protein content was determined using the Brachidonic-acid assay kit (Thermo Fisher Scientific, Cat# 23227). Equal loads of proteins were separated with the Mini-PROTEAN TGX system (Bio-Rad Laboratories) and transferred to polyvinylidene difluoride membrane (Bio-Rad Laboratories). Membranes were incubated with anti-FLAG antibody (1:400, Sigma Aldrich, Cat# F3165) overnight at 4℃. After incubation with horseradish peroxidase-conjugated secondary antibody (IgG detector, Takara Clontech, Cat# T7122A-1), the antigen was detected using chemiluminescence Western blotting detection reagents (Pierce ECL Western Blotting Substrate, Thermo Fisher Scientific Inc., Rockford, IL, the USA, Cat# 32106). The image was scanned with ChemiDoc (Bio-Rad Laboratories). Membranes were striped using stripping buffer (Restore Western Blotting Stripping Buffer, Thermo Fisher Scientific Inc., Cat# 21059) and re-probed using anti-beta actin (1:5000, Rabbit IgG, Cell Signaling Technology, Cat# 13E5) using the same steps that were used previously.

### Annexin V flowcytometry (FACS) assay

Apoptosis and cell death were analyzed using flowcytometry Annexin V assay as per our published protocol ^10^. In brief, following exposure to stress in 24 well plate, cells were collected by trypsinization using 0.25% Trypsin in EDTA and washed using Annexin V binding buffer (0.5% bovine serum albumin (BSA) + 0.2mM Calcium Chloride in Fluorobrite DMEM medium [Gibco, Cat# A18967-01]). Then, cells were incubated with Alexa Fluor 647 Annexin V antibody (Biolegend, Cat# 640943) and Propidium Idodie (Invitrogen, Cat# P3566) for 15 minutes. The samples were analyzed using FACS canto II. Statistical analysis (Student’s t-test) was conducted on SPSS v.20 (IBM Corp.). Statistical significance was considered when *p* was less than 0.5 and fold change of > 1.5 folds were observed.

### Real-time PCR analysis of gene expression (qPCR)

Seventy-two hours after plasmid transfection, cells were lysed in Qiazol and RNA extracted as above. cDNA was synthesized from 1000ng total RNA using iScript cDNA synthesis kit (Biorad, Cat# 1708891). Quantitative real-time PCR was performed on CFX96 (Bio-Rad) using GoTaq qPCR Master Mix (Promega Co., Madison, WI, USA). Primers used in this study were as follows:

Beta-actin: Forward: 5’GGAGATTACTGCCCTGGCTCCTA, Reverse: 5’GACTCATCGTACTCCTGCTTGCTG

Ribonuclease inhibitor 1 (RNH1): Forward: 5’GAGCTCAGCCTACGCACCAA, Reverse: 5’CACAGCCAGCTTCCGTCAA

ALKBH1: Forward: 5’TGGGTCACTCTGGGCTACCATTA, Reverse: 5’TCTGCTCGGAAACCCTGAAATC

## Acknowledgment

The authors would like to thank Dr Yasutoshi Akiyama for his insight and various comments made regarding this work. The authors would also like to thank Dr Peter C Dedon for his comments and insight into sequencing limitations that helped with the writing of the discussion section.

## Author contribution

**Rashad:** Conception and study design, administration, Funding acquisition, conducted all experiments, analyzed the data, and wrote the manuscript. **Niizuma:** Funding acquisition and administration. **Niizuma, Tominaga:** critically revised the manuscript. **All authors:** approved the final version of the manuscript.

## References

1 Advani, V. M. & Ivanov, P. Translational Control under Stress: Reshaping the Translatome. BioEssays : news and reviews in molecular, cellular and developmental biology 41, e1900009, doi:10.1002/bies.201900009 (2019).

2 Torrent, M., Chalancon, G., de Groot, N. S., Wuster, A. & Madan Babu, M. Cells alter their tRNA abundance to selectively regulate protein synthesis during stress conditions. Science signaling 11, doi:10.1126/scisignal.aat6409 (2018).

3 Goodarzi, H. et al. Endogenous tRNA-Derived Fragments Suppress Breast Cancer Progression via YBX1 Displacement. Cell 161, 790–802, doi:10.1016/j.cell.2015.02.053 (2015).

4 Liu, F. et al. ALKBH1-Mediated tRNA Demethylation Regulates Translation. Cell 167, 816–828 e816, doi:10.1016/j.cell.2016.09.038 (2016).

5 Pollo-Oliveira, L. et al. Loss of Elongator- and KEOPS-Dependent tRNA Modifications Leads to Severe Growth Phenotypes and Protein Aggregation in Yeast. Biomolecules 10, doi:10.3390/biom10020322 (2020).

6 Rashad, S., Niizuma, K. & Tominaga, T. tRNA cleavage: a new insight. Neural regeneration research 15, 47–52, doi:10.4103/1673-5374.264447 (2020).

7 Ivanov, P., Emara, M. M., Villen, J., Gygi, S. P. & Anderson, P. Angiogenin-induced tRNA fragments inhibit translation initiation. Molecular cell 43, 613–623, doi:10.1016/j.molcel.2011.06.022 (2011).

8 Goncalves, K. A. et al. Angiogenin Promotes Hematopoietic Regeneration by Dichotomously Regulating Quiescence of Stem and Progenitor Cells. Cell 166, 894–906, doi:10.1016/j.cell.2016.06.042 (2016).

9 Sato, K., Rashad, S., Niizuma, K. & Tominaga, T. Stress induced tRNA halves (tiRNAs) as biomarkers for stroke and stroke therapy; Pre-clinical study. Neuroscience, doi:10.1016/j.neuroscience.2020.03.018 (2020).

10 Elkordy, A. et al. tiRNAs as a novel biomarker for cell damage assessment in in vitro ischemia-reperfusion model in rat neuronal PC12 cells. Brain research 1714, 8–17, doi:10.1016/j.brainres.2019.02.019 (2019).

11 Dhahbi, J. M. et al. 5' tRNA halves are present as abundant complexes in serum, concentrated in blood cells, and modulated by aging and calorie restriction. BMC genomics 14, 298, doi:10.1186/1471-2164-14-298 (2013).

12 Elkordy, A. et al. Stress-induced tRNA cleavage and tiRNA generation in rat neuronal PC12 cells. J Neurochem 146, 560–569, doi:10.1111/jnc.14321 (2018).

13 Yamasaki, S., Ivanov, P., Hu, G. F. & Anderson, P. Angiogenin cleaves tRNA and promotes stress-induced translational repression. The Journal of cell biology 185, 35–42, doi:10.1083/jcb.200811106 (2009).

14 Krishna, S. et al. Dynamic expression of tRNA-derived small RNAs define cellular states. EMBO Rep, doi:10.15252/embr.201947789 (2019).

15 Su, Z., Kuscu, C., Malik, A., Shibata, E. & Dutta, A. Angiogenin generates specific stress-induced tRNA halves and is not involved in tRF-3-mediated gene silencing. The Journal of biological chemistry 294, 16930–16941, doi:10.1074/jbc.RA119.009272 (2019).

16 Akiyama, Y. et al. Multiple ribonuclease A family members cleave transfer RNAs in response to stress. bioRxiv, 811174, doi:10.1101/811174 (2019).

17 Rashad, S. et al. The stress specific impact of ALKBH1 on tRNA cleavage and tiRNA generation. RNA biology, doi:10.1080/15476286.2020.1779492 (2020).

18 Ivanov, P. et al. G-quadruplex structures contribute to the neuroprotective effects of angiogenin-induced tRNA fragments. Proceedings of the National Academy of Sciences of the United States of America 111, 18201–18206, doi:10.1073/pnas.1407361111 (2014).

19 Emara, M. M. et al. Angiogenin-induced tRNA-derived stress-induced RNAs promote stress-induced stress granule assembly. J Biol Chem 285, 10959–10968, doi:10.1074/jbc.M109.077560 (2010).

20 Kawarada, L. et al. ALKBH1 is an RNA dioxygenase responsible for cytoplasmic and mitochondrial tRNA modifications. Nucleic acids research 45, 7401–7415, doi:10.1093/nar/gkx354 (2017).

21 Chen, Z. et al. Transfer RNA demethylase ALKBH3 promotes cancer progression via induction of tRNA-derived small RNAs. Nucleic acids research 47, 2533–2545, doi:10.1093/nar/gky1250 (2019).

22 Tuorto, F. et al. RNA cytosine methylation by Dnmt2 and NSun2 promotes tRNA stability and protein synthesis. Nature structural & molecular biology 19, 900–905, doi:10.1038/nsmb.2357 (2012).

23 Rashad, S. et al. The stress specific impact of ALKBH1 on tRNA cleavage and tiRNA generation. RNA biology, 1–12, doi:10.1080/15476286.2020.1779492 (2020).

24 Faheem, H. et al. Neuroprotective effects of minocycline and progesterone on white matter injury after focal cerebral ischemia. Journal of clinical neuroscience : official journal of the Neurosurgical Society of Australasia 64, 206–213, doi:10.1016/j.jocn.2019.04.012 (2019).

25 Li, Q. et al. tRNA-Derived Small Non-Coding RNAs in Response to Ischemia Inhibit Angiogenesis. Sci Rep 6, 20850, doi:10.1038/srep20850 (2016).

26 Zhou, R., Yazdi, A. S., Menu, P. & Tschopp, J. A role for mitochondria in NLRP3 inflammasome activation. Nature 469, 221–225, doi:10.1038/nature09663 (2011).

27 Lyons, S. M., Fay, M. M., Akiyama, Y., Anderson, P. J. & Ivanov, P. RNA biology of angiogenin: Current state and perspectives. RNA biology 14, 171–178, doi:10.1080/15476286.2016.1272746 (2017).

28 Thomas, S. P., Hoang, T. T., Ressler, V. T. & Raines, R. T. Human angiogenin is a potent cytotoxin in the absence of ribonuclease inhibitor. Rna 24, 1018–1027, doi:10.1261/rna.065516.117 (2018).

29 Pizzo, E. et al. Ribonuclease/angiogenin inhibitor 1 regulates stress-induced subcellular localization of angiogenin to control growth and survival. J Cell Sci 126, 4308–4319, doi:10.1242/jcs.134551 (2013).

30 Saikia, M. et al. Genome-wide identification and quantitative analysis of cleaved tRNA fragments induced by cellular stress. J Biol Chem 287, 42708–42725, doi:10.1074/jbc.M112.371799 (2012).

31 Furia, A. et al. The ribonuclease/angiogenin inhibitor is also present in mitochondria and nuclei. FEBS letters 585, 613–617, doi:10.1016/j.febslet.2011.01.034 (2011).

32 Ougland, R. et al. ALKBH1 is a histone H2A dioxygenase involved in neural differentiation. Stem cells (Dayton, Ohio) 30, 2672–2682, doi:10.1002/stem.1228 (2012).

33 Hu, J. et al. Sequencing-based quantitative mapping of the cellular small RNA landscape. bioRxiv, 841130, doi:10.1101/841130 (2019).

34 Akiyama, Y., Kharel, P., Abe, T., Anderson, P. & Ivanov, P. Isolation and initial structure-functional characterization of endogenous tRNA-derived stress-induced RNAs. RNA biology, 1–9, doi:10.1080/15476286.2020.1732702 (2020).

35 Blázquez, M., Fominaya, J. M. & Hofsteenge, J. Oxidation of sulfhydryl groups of ribonuclease inhibitor in epithelial cells is sufficient for its intracellular degradation. The Journal of biological chemistry 271, 18638–18642, doi:10.1074/jbc.271.31.18638 (1996).

36 Schneider, C. A., Rasband, W. S. & Eliceiri, K. W. NIH Image to ImageJ: 25 years of image analysis. Nature methods 9, 671–675, doi:10.1038/nmeth.2089 (2012).

