## Supplementary figures for "The cell and stress-specific canonical and non-canonical tRNA cleavage"

### Supplementary data

**Supplementary Figure 1: Evaluation of the impact of arsenite (AS; 1mM) or antimycin A (A.M; 200μg/ml) on cell death using Annexin V flowcytometry and tRNA cleavage using SYBR gold staining in PC12 cells (A) and B35 cells (B). A:** As caused cell death and tRNA cleavage in PC12 cells while A.M failed to achieve either. **B:** As and A.M caused cell death and tRNA cleavage in B35 cells with A.M causing a more robust tRNA cleavage.

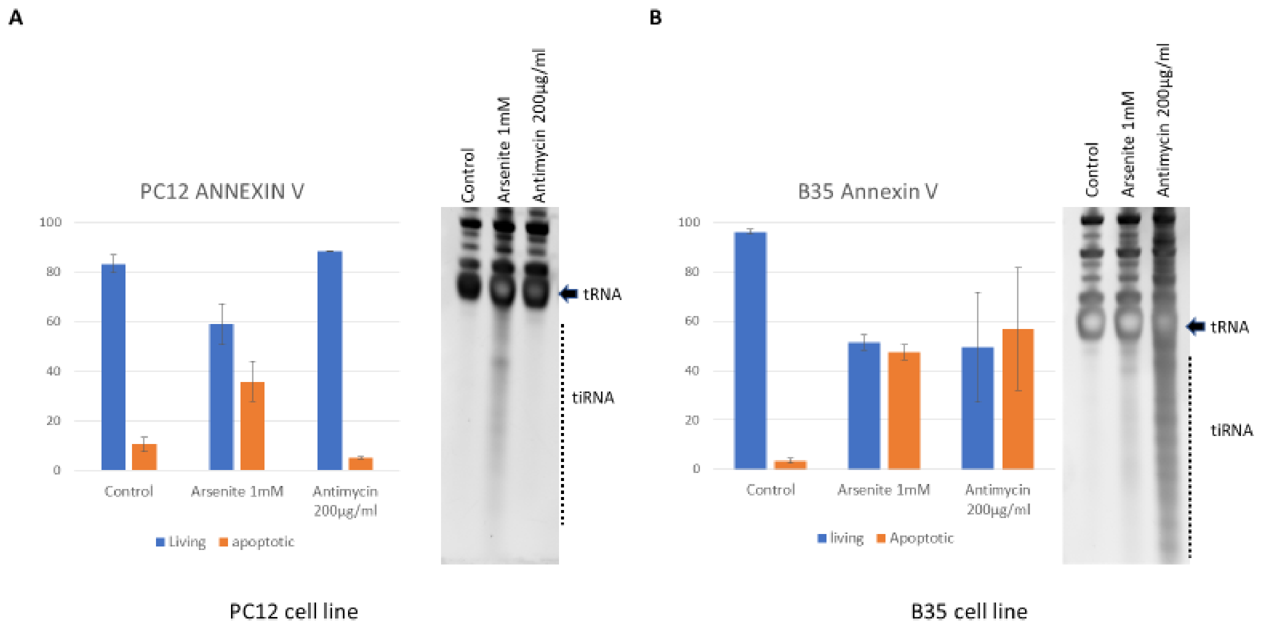

**Supplementary Figure 2:** Shorter exposure images for overexposed blots in Figure 2

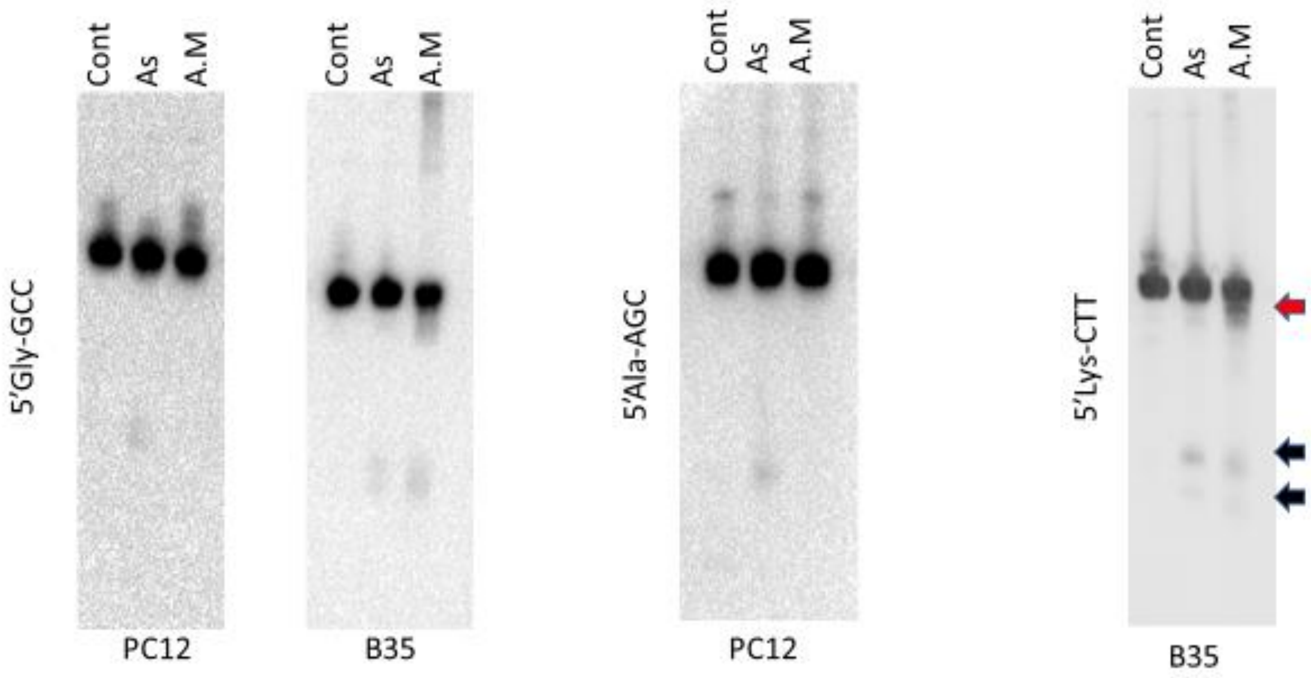

**Supplementary Figure 3: Impact of hAng and RNH1 on gross tRNA cleavage: A:** western blotting confirmation of Flag labeled human angiogenin expression in PC12. **B:** qPCR analysis of RNH1 gene expression following plasmid transfection in PC12 reveals >80 upregulation of RNH1. **C:** SYBR gold staining after hAng overexpression and stress shows that hAng overexpression increased tRNA cleavage after As stress only and had not impact on A.M stress. **D:** RNH1 overexpression did not suppress tRNA cleavage after As stress. Arrow: tRNA, dotted line: tiRNA.

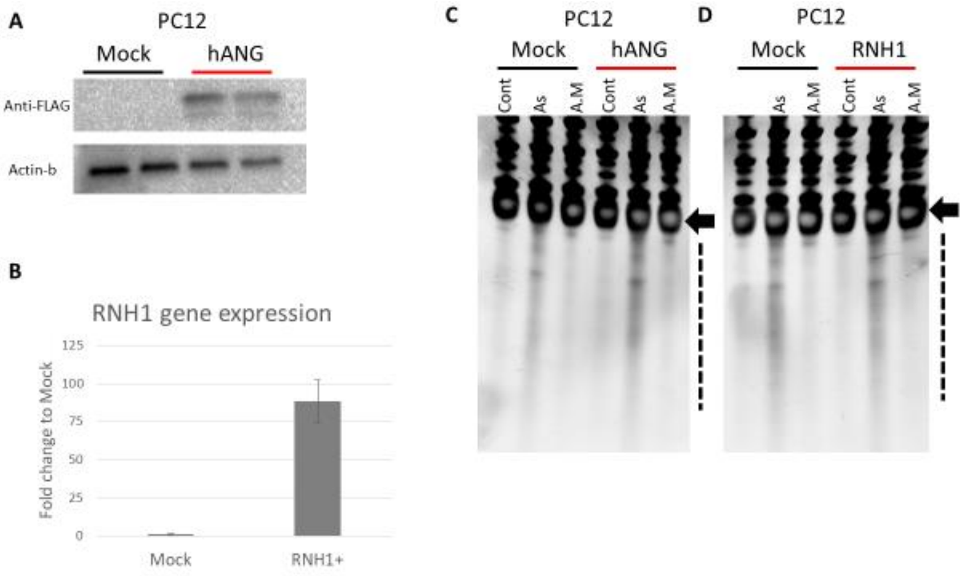

**Supplementary Figure 4:** Shorter exposure images for the overexposed blots in Figures 4 and 5 and 5s rRNA loading control blot

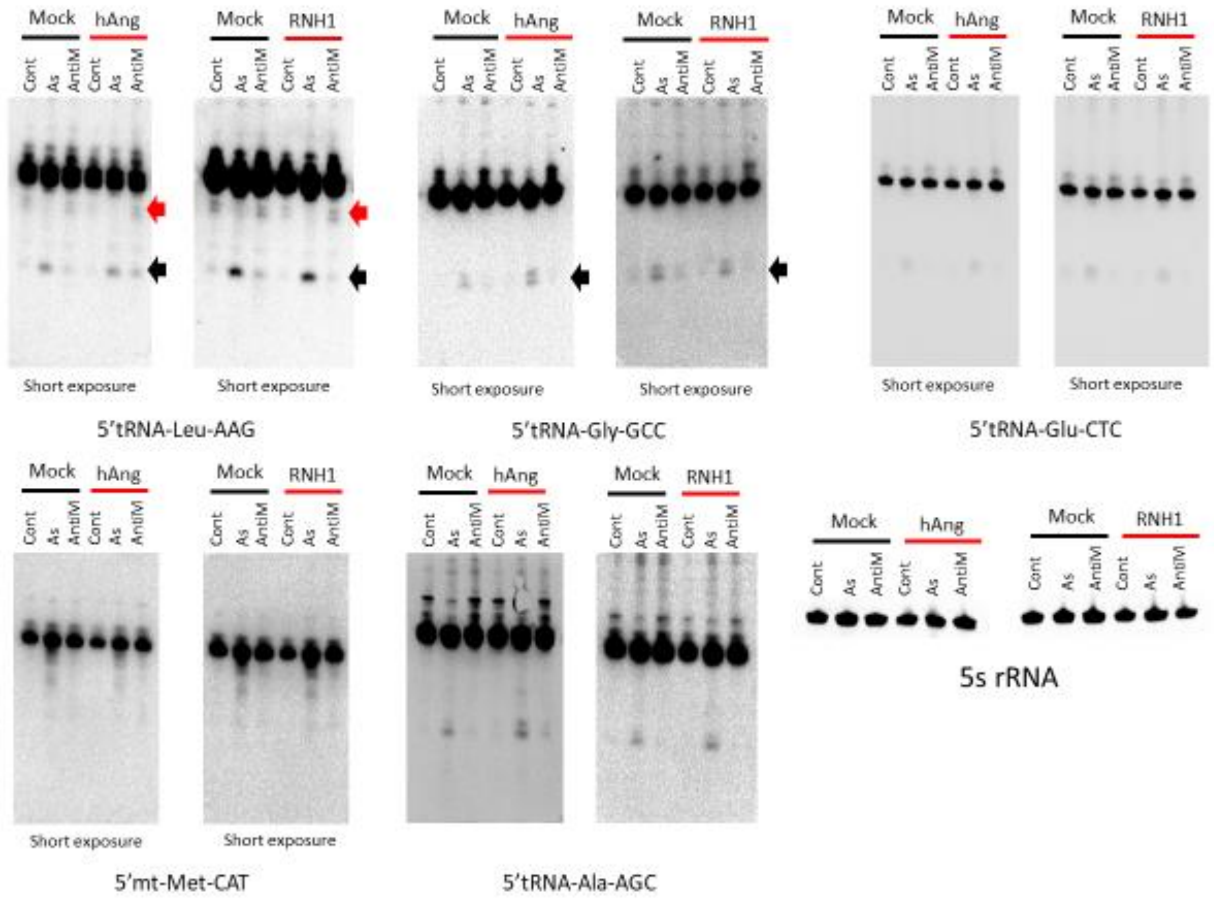

**Supplementary Figure 5: Confirmation of Alkbh1 overexpression** after plasmid transfection of PC12 reveals >400 folds upregulation in gene expression.

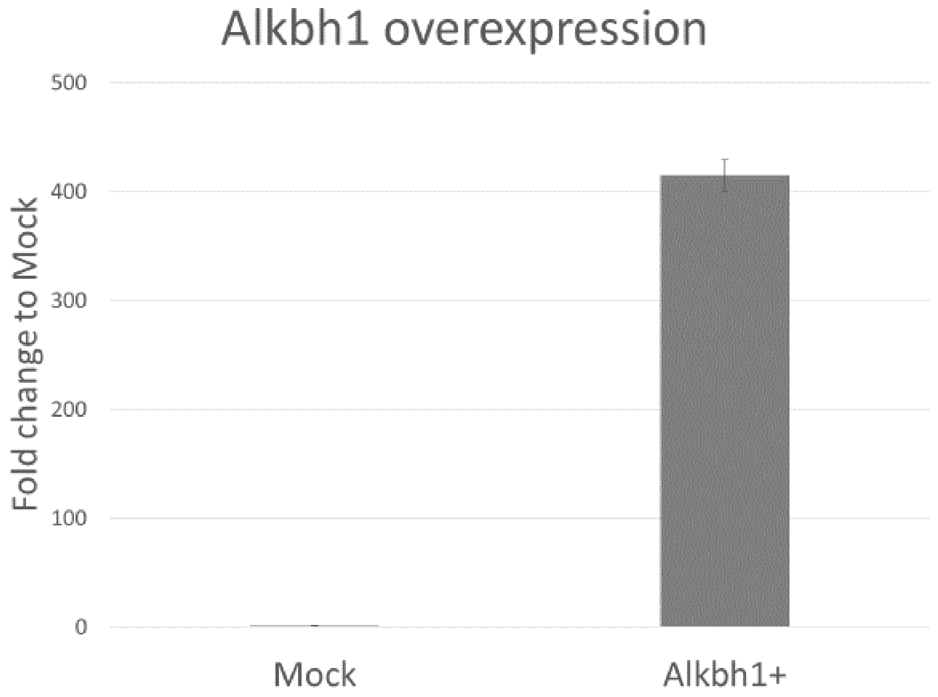

**Supplementary Figure 6: Shorter exposure for blots in Figure 6 plus 5s rRNA presented as loading control**

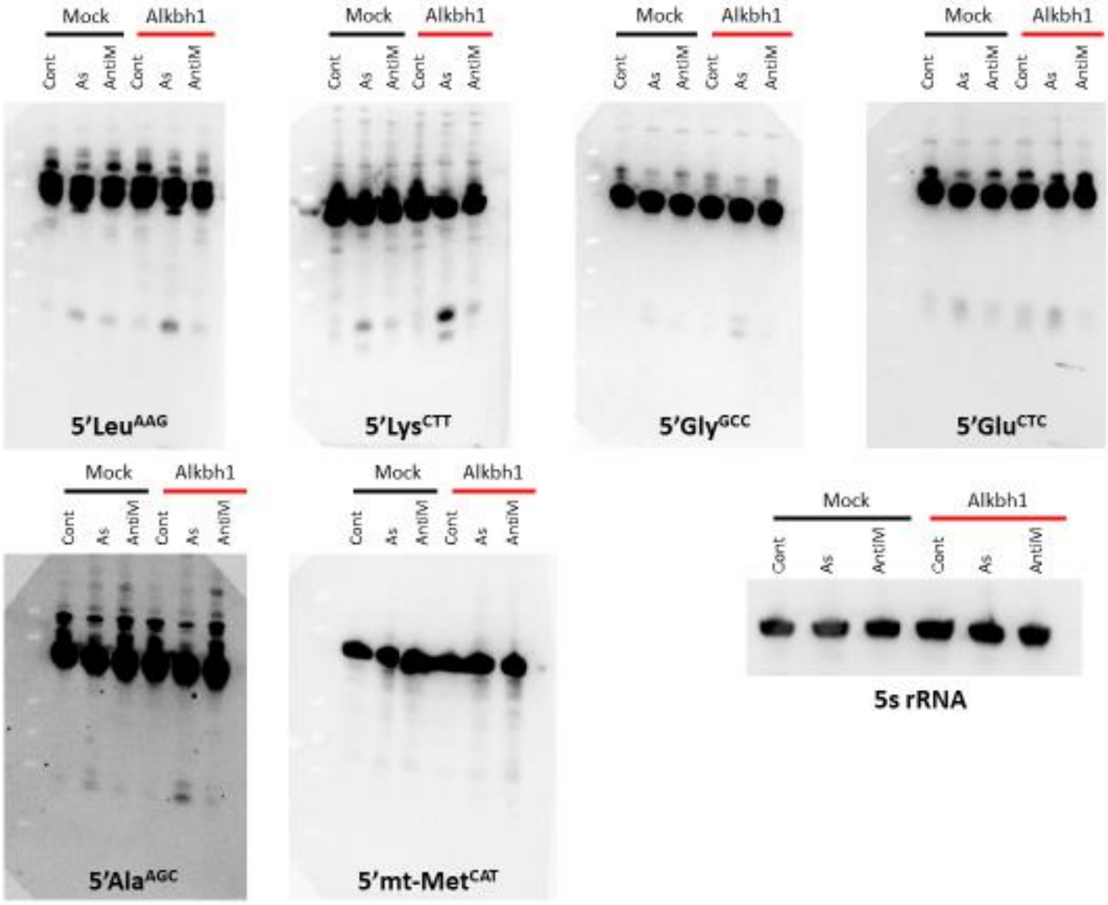
